# Deep phylo-taxono-genomics (DEEPT genomics) reveals misclassification of *Xanthomonas* species complexes into *Xylella, Stenotrophomonas* and *Pseudoxanthomonas*

**DOI:** 10.1101/2020.02.04.933507

**Authors:** Kanika Bansal, Sanjeet Kumar, Amandeep Kaur, Shikha Sharma, Prashant Patil, Prabhu B. Patil

## Abstract

Genus *Xanthomonas* encompasses specialized group of phytopathogenic bacteria with genera *Xylella, Stenotrophomonas* and *Pseudoxanthomonas* being its closest relatives. While species of genera *Xanthomonas* and *Xylella* are known as serious phytopathogens, members of other two genera are found in diverse habitats with metabolic versatility of biotechnological importance. Few species of *Stenotrophomonas* are multidrug resistant opportunistic nosocomial pathogens. In the present study, we report genomic resource of genus *Pseudoxanthomonas* and further in-depth comparative studies with publically available genome resources of other three genera. Surprisingly, based on deep phylo-taxono-genomic or DEEPT criteria, all the four genera were found to be one genus and hence synonyms of *Xanthomonas*. The members of *Pseudoxanthomonas* are more diverse and ancestral and rest forming two mega species groups (MSGs) i.e. *Xanthomonas Xylella* (XX-MSG) and *Stenotrophomonas* (S-MSG). Within XX-MSG, there are 3 species complexes i.e., *X. citri* complex (Xcc) member species are primarily pathogenic to dicots, *X. hyacinthi* complex (Xhc) member species are primarily pathogenic to monocots and *Xanthomonas* (*Xylella*) *fastidosa* complex (Xfc) with diverse phytopathogenic potential. Interestingly, *Xylella* seems to be a variant *Xanthomonas* lineage or species complex that is sandwiched between Xcc and Xhc. Like *Xylella*, within S-MSG, we find a species complex of clinical origin *Xanthomonas* (*Stenotrophomonas*) *maltophilia* complex (Xmc). Comparative studies revealed selection and role of xanthomonadin pigment and xanthan gum in emergence of plant pathogenic XX-MSG. Pan genome analysis also revealed large set of unique genes with particular functions suited for plant/animal lifestyle responsible for emergence of variant Xfc and Xmc species complexes. Overall, our systematic and large scale genera based study has allowed us to understand the origin and to clarify the taxonomic breadth of genus of high importance in agriculture, medicine and industry. Such DEEPT genomics studies are also way forward to identify right markers or functions for diagnosis and drug development of any pathogenic bacteria.

**Repositories:** Genome Submission Accession Number:

MWIP00000000, PDWO00000000, PDWN00000000, PDWT00000000, PDWS00000000, PDWW00000000, PDWU00000000, PDWR00000000, PDWL00000000, PDWQ00000000, PDWM00000000, PDWP00000000, PDWV00000000, PDWK00000000 and QOVG00000000

## Introduction

*Xanthomonadaceae* (*Lysobacteraceae*) consists of genera from diverse ecological niches following different lifestyles. Sequence based studies using conserved signature indels (CSI), housekeeping genes and phylogenomics of type species/ representative members of its genera and families revealed that *Xanthomonas, Stenotrophomonas, Xylella* and *Pseudoxanthomonas* are closely related forming a phylogroup (referred here as XSXP phylogroup) (Naushad and Gupta 2013, Kumar, Bansal et al. 2019). Amongst XSXP, both *Xanthomonas* and *Xylella* are known to have characteristic plant pathogenic lifestyle (Leyns, De Cleene et al. 1984, Hopkins 1989). Whereas, *Stenotrophomonas* (http://www.bacterio.net/stenotrophomonas.html) and *Pseudoxanthomonas* (http://www.bacterio.net/pseudoxanthomonas.html) are versatile group of bacteria found in diverse environments including water, soil, contaminated sites etc. and are reported to have biotechnological significance (Ryan, Monchy et al. 2009, Patel, Cheturvedula et al. 2012, Mahbub, Krishnan et al. 2016, Mohapatra, Sar et al. 2018). Moreover, members of *Stenotrophomonas maltophilia* complex are listed by WHO as priority multidrug resistant pathogens (http://www.who.int/drugresistance/AMR_Importance/en/).

According to classical taxonomy there are various conflicts within XSXP phylogroup. For instance, type species of genus *Stenotrophomonas* i.e. *S. maltophilia* was also classified as *Xanthomonas maltophilia* (Palleroni and Bradbury 1993). Similarly, type species of the genus *Pseudoxanthomonas* i.e. *P. broegbernensis* (Finkmann, Altendorf et al. 2000) was separated from *Xanthomonas* and *Stenotrophomonas* based on the absence of the fatty acid 3-hydroxy-iso-tridecanoic acid and by their ability to reduce the nitrite but not nitrate to N_2_O and from genus *Xylella* by the presence of branched-chain fatty acids (Yang, Vauterin et al. 1993, Assih, Ouattara et al. 2002, Thierry, Macarie et al. 2004). Further, genus *Xylella* is having highly reduced genome and GC content in the XSXP phylogroup, hence, taxonomic and phylogenetic status of *Xylella* needs critical examination (Wells, Raju et al. 1987, Finkmann, Altendorf et al. 2000, Simpson, Reinach et al. 2000). Considering importance of members of these genera in agriculture, medicine and industry there is a need to carry out systematic and large scale genome based studies of XSXP genera.

Advent of cost-effective and high-throughput genomics era is transforming the way we understand the relationship of bacteria (Garrity 2016). Whole genome information will not only enable us to understand the emergence of new lineages, strains or clones but also establish identity along with relationship of a bacterium at the level of species and genera (Konstantinidis and Tiedje 2005, Konstantinidis and Tiedje 2005).While for species delineation average nucleotide identity (ANI) and digital DNA-DNA hybridization (dDDH) are highly accurate with 96% and 70% cut-offs, respectively (Richter and Rosselló-Móra 2009, Auch, von Jan et al. 2010). New criteria are also becoming available for delineation of members at the genus level. Apart from phylogenetic trees based on large set of genomic markers and core genome, there are criteria like average amino acid identity (AAI) and percentage of conserved protein (POCP) that have been proposed for genus delineation with 60-80% and 50% cut-offs, respectively (Konstantinidis and Tiedje 2005, Qin, Xie et al. 2014). POCP is affected by genome size and hence cannot be used for the species undergoing extreme genome reduction like *Xylella* (Qin, Xie et al. 2014, Hayashi Sant’Anna, Bach et al. 2019). However, core genome based trees and AAI are not affected by genome reduction and can be employed to establish identity and relationship of genera irrespective of genome size and or GC content (Konstantinidis and Tiedje 2005, Hayashi Sant’Anna, Bach et al. 2019, Indu, Ch et al. 2019).

Due to economic importance, genome resource of type strains of genus *Xanthomonas* (https://www.ncbi.nlm.nih.gov/genome/?term=xanthomonas) and *Xylella* (https://www.ncbi.nlm.nih.gov/genome/?term=xylella) is publically available. In an earlier study from our group, we reported genomes and taxonogenomic study of type strains of the genus *Stenotrophomonas* (Patil, Midha et al. 2016). But, genomic resource of type strains of *Pseudoxanthomonas* is warranted for large scale genome level evolutionary study of these genera. In this study, we have generated genomic resource of type strains of the genus *Pseudoxanthomonas* and utilized available genomic resource of other three genera to carry out deep genome based phylogenetic, taxonomic and evolutionary studies. Present study reveals that all these four genera actually belong to one genus and hence synonym of genus *Xanthomonas*. Further, phylogenomic investigation revealed that members of the genus *Pseudoxanthomonas* form diverse backbone to other three genera. Genus *Xylella* is phylogenomically closer to *Xanthomonas* than *Stenotrophomonas* and is in fact a variant intermediate or variant lineage or species complex that is sandwiched between two plant pathogenic *Xanthomonas* species complexes associated with dicots and monocots. Interestingly, our study revealed species complex associated with clinical species is also a variant lineage or species complex within the genus *Stenotrophomonas.*

Comparative gene-content studies revealed importance of a pigment and exopolysaccharide in origin of *Xanthomonas Xylella* MSG and large set of unique genes in emergence of variant species complexes whose members have animal/plant bimodal lifestyle. The genomic resource, unique gene sets and counterintuitive findings from this deep phylo-taxono-genomic or DEEPT genomic study of XSXP have major implications in understanding, classification and management of pathogenic species/species complexes of importance in agriculture and medicine. At the same time, it will also help in systematic exploitation of the species of biotechnological importance.

## Results

### Genomic resource of genus *Pseudoxanthomonas*

Genome sequencing of 15 *Pseudoxanthomonas* type strains was carried out in-house (Pl. see methods) and raw reads were assembled *de novo* resulting in draft genomes with minimum contig size of 500 bp. The range of genome size, coverage and N50 were 3.03 Mb to 4.6 Mb, 32x to 270x and 39.9 Kb to 610.8 Kb respectively. Average GC% of the genomes was 68.13%. A detailed genome feature and assembly statistics are provided in table 1.

**Table 1:** Genome assembly statistics of species of the genus *Pseudoxanthomonas*.

| Strain name | Genome Size (bp) | Fold | #Contigs | N50 (bp) | %GC | Completeness/<br>Contamination | # CDS | tRNA +<br>rRNA | Accession Number |
| --- | --- | --- | --- | --- | --- | --- | --- | --- | --- |
| *P. broegbernensis* DSM 12573<sup>T</sup> | 3,547,767 | 267x | 157 | 175278 | 70.6 | 99.66/1.74 | 3024 | 54+3 | MWIP00000000 |
| *P. kaohsiungensis* DSM 17583<sup>T</sup> | 3774556 | 269x | 100 | 199653 | 69.68 | 99.66/1.42 | 3420 | 52+3 | PDWO00000000 |
| *P. daejeonensis* DSM 17801<sup>T</sup> | 3563566 | 131x | 38 | 227294 | 68.89 | 99.66/0.11 | 3143 | 58+3 | PDWN00000000 |
| *P. yeongjuensis* DSM 18204<sup>T</sup> | 3937688 | 137x | 28 | 610842 | 65.11 | 99.95/0.76 | 3389 | 53+3 | PDWT00000000 |
| *P. sacheonensis* DSM 19373<sup>T</sup> | 4036514 | 101x | 44 | 165077 | 64.3 | 100/0.91 | 3582 | 50+3 | PDWS00000000 |
| *P. japonensis* DSM 17109<sup>T</sup> | 4075711 | 111x | 74 | 108688 | 67.3 | 99.95/0.34 | 3673 | 50+3 | PDWW00000000 |
| *P. wuyuanensis* DSM 100640<sup>T</sup> | 4686433 | 112x | 86 | 149132 | 65.75 | 100/1.08 | 4003 | 51+6 | PDWU00000000 |
| *P. sangjuensis* DSM 28345<sup>T</sup> | 3289016 | 91x | 76 | 105206 | 68.68 | 99.95/0.46 | 2908 | 51+3 | PDWR00000000 |
| *P. jiangsuensis* DSM 22398<sup>T</sup> | 3790571 | 171x | 154 | 121587 | 70.35 | 99.31/1.03 | 3389 | 50+3 | PDWL00000000 |
| *P. kalamensis* DSM 18571<sup>T</sup> | 3034522 | 239x | 162 | 478688 | 65.88 | 99.84/1.50 | 2697 | 48+3 | PDWQ00000000 |
| *P. koreensis* KCTC12208<sup>T</sup> | 3049736 | 74x | 93 | 60801 | 70.17 | 99.48/0.04 | 2681 | 52+3 | PDWM00000000 |
| *P. suwonensis* DSM 17175<sup>T</sup> | 3428455 | 65x | 66 | 104754 | 70.36 | 99.66/0.34 | 3071 | 55+3 | PDWP00000000 |
| *P. mexicana* DSM17121<sup>T</sup> | 3965467 | 32x | 115 | 63754 | 67.4 | 99.49/1.71 | 3660 | 54+6 | PDWV00000000 |
| *P. taiwanensis* DSM22914<sup>T</sup> | 3043352 | 69x | 160 | 39985 | 72.08 | 99.16/0.34 | 2729 | 56+3 | PDWK00000000 |
| *P. gei* KCTC32298<sup>T</sup> | 3431103 | 173x | 50 | 332530 | 65.46 | 99.95/0.41 | 3,122 | 45+4 | QOVG00000000 |

### Genomic features of XSXP phylogroup

Genomic features of type strains of the XSXP phylogroup are summarized in table 2. Genome size of *Xanthomonas* is around 5 Mb and around 3-5 Mb for *Pseudoxanthomonas* and *Stenotrophomonas*, whereas, genome size of *Xylella* is around 2.5 Mb. This reduction in genome size is reflected in number of coding sequences as it was in range of 3000 to 4000 for *Xanthomonas, Stenotrophomonas* and *Pseudoxanthomonas*, but around 2100 for *Xylella.* Furthermore, average GC content for all was around 65% except for *Xylella*, which was having GC content of around 51%. In spite of violent genome changes, *Xylella* is displaying comparable number of tRNAs i.e. 48.

**Table 2:** Metadata of the strains used in the present study. Following data was obtained from list of prokaryotic names with standing in nomenclature (http://www.bacterio.net/) and NCBI (https://www.ncbi.nlm.nih.gov/). Star marked columns are obtained in the present study.

| Strain | Genome size | CDS | Genome status | Source/Host | Host taxonomy | Lifestyle | Location | Isolation year | GC % | tRNA | Completeness*/Contaminations* |
| --- | --- | --- | --- | --- | --- | --- | --- | --- | --- | --- | --- |
| *Xanthomonas vasicola* NCPPB 2417<sup>T</sup> | 4.9 | 3960 | Draft | *Sorghum vulgare* | Tracheophytes, Angiosperms, Monocots, Commelinids, Poales, Poaceae | Pathogenic | New Zealand | 1969 | 63.3 | 51 | 100/0.07 |
| *Xanthomonas oryzae* ATCC 35933<sup>T</sup> | 4.2 | 3385 | Draft | *Oryza sativa* (leaf blight and wilt) | Tracheophytes, Angiosperms, Monocots, Commelinids, Poales, Poaceae | Pathogenic | - | - | 63.9 | 49 | 100/0 |
| *Xanthomonas euvesicatoria* 85-10 | 5.1 | 4416 | Complete | Infected pepper plant | Tracheophytes, Angiosperms, Eudicots, Asterids, Solanales, Solanaceae | Pathogenic | - | - | 64.7 | 54 | 99.64/0 |
| *Xanthomonas alfalfae* LMG 495<sup>T</sup> | 5.0 | 3937 | Draft | *Medicago sativa* | Tracheophytes, Angiosperms, Eudicots, Rosids, Fabales, Fabaceae | Pathogenic | Belgium: Merelbeke | 2014 | 64.8 | 44 | 98.79/2.99 |
| *Xanthomonas perforans* CFBP 7293<sup>T</sup> | 5.0 | 4206 | Draft | Leaves of *Solanum lycopersicum* | Tracheophytes, Angiosperms, Eudicots, Asterids, Solanales, Solanaceae | Pathogenic | USA: Florida | - | 64.9 | 54 | 99.64/0.2 |
| *Xanthomonas phaseoli* CFBP 412 | 5.0 | 4175 | Complete | *Phaseolus vulgaris* | Tracheophytes, Angiosperms, Eudicots, Rosids, Fabales, Fabaceae | Pathogenic | USA | - | 65 | 54 | 99.64/0.24 |
| *Xanthomonas citri* LMG 9322<sup>T</sup> | 5.1 | 435<br>5 | Draft | *Citrus aurantifolia* | Tracheophytes,<br>Angiosperms,<br>Eudicots, Rosids,<br>Sapindales, Rutaceae | Pathogenic | USA: Florida | 1915 | 64.6 | 50 | 98.19/1.09 |
| *Xanthomonas axonopodis* DSM 3585<sup>T</sup> | 4.4 | 326<br>3 | Draft | *Axonopus scoparius* | Tracheophytes,<br>Angiosperms,<br>Monocots,<br>Commelinids,<br>Poales, Poaceae | Pathogenic | Colombia | 1949 | 64.5 | 50 | 99.52/0.12 |
| *Xanthomonas prunicola* CFBP 8353<sup>T</sup> | 5.3 | 420<br>8 | Draft | *Prunus persica* var.<br>nectarina | Tracheophytes,<br>Angiosperms,<br>Eudicots, Rosids,<br>Rosales, Rosaceae | Pathogenic | Spain:<br>Murcia,<br>Abaran | 2015 | 64 | 53 | 100/0 |
| *Xanthomonas bromi* CFBP 1976<sup>T</sup> | 4.9 | 384<br>5 | Draft | *Bromus carinatus* | Tracheophytes,<br>Angiosperms,<br>Monocots,<br>Commelinids,<br>Poales, Poaceae | Pathogenic | France | 1980 | 64 | 53 | 100/0 |
| *Xanthomonas nasturtii* WHRI 8853<sup>T</sup> | 4.8 | 398<br>1 | Draft | Watercress | Tracheophytes,<br>Angiosperms,<br>Eudicots, Rosids,<br>Brassicales,<br>Brassicaceae | Pathogenic | USA: Florida | 2014 | 64.5 | 56 | 100/0.36 |
| *Xanthomonas cynarae* CFBP 4188<sup>T</sup> | 5.0 | 410<br>2 | Draft | *Cynara scolymus* | Tracheophytes,<br>Angiosperms,<br>Eudicots, Asterids,<br>Asterales, Asteraceae | Pathogenic | France:<br>Bretagne | 1996 | 63.7 | 51 | 100/0.41 |
| *Xanthomonas gardneri* ICMP 7383 | 5.3 | 452<br>3 | Complete | Tomato field | Tracheophytes,<br>Angiosperms,<br>Eudicots, Asterids,<br>Solanales,<br>Solanaceae | Pathogenic | New Zealand | 1980 | 63.5 | 53 | 99.64/0.68 |
| *Xanthomonas hortorum* WHRI 7744<sup>T</sup> | 5.5 | 442<br>7 | Draft | *Hedera helix* | Tracheophytes,<br>Angiosperms,<br>Eudicots, Asterids,<br>Apiales, Araliaceae | Pathogenic | USA | 1961 | 63.6 | 54 | 100/1.03 |
| *Xanthomonas populi* CFBP 1817<sup>T</sup> | 4.3 | 322<br>2 | Draft | *Populus x canadensis* cv. regenerata) | Tracheophytes, Angiosperms, Eudicots, Rosids, Malpighiales, Salicaceae | Pathogenic | Oise, Noyon | 1957 | 63.3 | 48 | 99.64/0.14 |
| *Xanthomonas arboricola* CFBP 2528<sup>T</sup> | 5.0 | 412<br>4 | Draft | Twig from a walnut tree | Tracheophytes, Angiosperms, Eudicots, Asterids, Fagales, Juglandaceae | Pathogenic | New Zealand | 1956 | 65.5 | 51 | 100/0.41 |
| *Xanthomonas fragariae* PD 885 | 4.2 | 326<br>3 | Complete | Diseased strawberry | Tracheophytes, Angiosperms, Eudicots, Rosids, Rosales, Rosaceae | Pathogenic | - | - | 62.3 | 52 | 99.64/0.36 |
| *Xanthomonas pisi* DSM 18956 | 4.2 | 338<br>5 | Draft | *Pisum sativum*; Leaf spot | Tracheophytes, Angiosperms, Eudicots, Rosids, Fabales, Fabaceae | Pathogenic | Japan | 1997 | 64.7 | 43 | 80.23/0 |
| *Xanthomonas dyei* CFBP 7245<sup>T</sup> | 5.3 | 427<br>8 | Draft | *Metrosideros excelsa*; Angular leaf lesion | Tracheophytes, Angiosperms, Eudicots, Rosids, Myrtales, Myrtaceae | Pathogenic | Bay of Plenty Omahanui | 1993 | 64.3 | 51 | 100/0.36 |
| *Xanthomonas vesicatoria* LMG 911<sup>T</sup> | 5.1 | 434<br>3 | Complete | *Lycopersicon esculentum* | Tracheophytes, Angiosperms, Eudicots, Asterids, Solanales, Solanaceae | Pathogenic | New Zealand | 1955 | 64.3 | 55 | 100/0.36 |
| *Xanthomonas melonis* CFBP 4644 | 4.6 | 387<br>2 | Draft | *Cucumis melo* | Tracheophytes, Angiosperms, Eudicots, Rosids, Cucurbitales, Cucurbitaceae | Pathogenic | Brazil | 1974 | 66.1 | 52 | 99.64/0.18 |
| *Xanthomonas cucurbitae* CFBP 2542<sup>T</sup> | 4.4 | 355<br>3 | Draft | *Cucurbita maxima* | Tracheophytes, Angiosperms, Eudicots, Rosids, Cucurbitales, Cucurbitaceae | Pathogenic | New Zealand | 1968 | 65.4 | 51 | 100/0.36 |
| *Xanthomonas floridensis* WHRI 8848<sup>T</sup> | 5.2 | 428<br>6 | Draft | Watercress | Tracheophytes,<br>Angiosperms,<br>Eudicots, Rosids,<br>Brassicales,<br>Brassicaceae | Pathogenic | USA: Florida | 2014 | 65.4 | 52 | 100/0.36 |
| *Xanthomonas cassavae* CFBP 4642<sup>T</sup> | 5.2 | 421<br>8 | Draft | *Manihot esculenta* | Tracheophytes,<br>Angiosperms,<br>Eudicots, Rosids,<br>Malpighiales,<br>Euphorbiaceae | Pathogenic | Malawi | 1951 | 65.2 | 51 | 99.64/0.93 |
| *Xanthomonas codiae* CFBP 4690 | 5.0 | 412<br>5 | Draft | *Codiaeum variegatum* | Tracheophytes,<br>Angiosperms,<br>Eudicots, Rosids,<br>Malpighiales,<br>Euphorbiaceae | Pathogenic | USA: Florida | 1987 | 66 | 53 | 99.28/0 |
| *Xanthomonas campestris* pv. *campestris* ATCC 33913<sup>T</sup> | 5.0 | 417<br>9 | Complete | Plant (*Brassica oleracea* var. *gemmifera*) | Tracheophytes,<br>Angiosperms, Eudicots, Rosids,<br>Brassicales,<br>Brassicaceae | Pathogenic | - | - | 65.1 | 53 | 99.64/0 |
| *Xanthomonas maliensis* LMG 27592<sup>T</sup> | 5.2 | 419<br>7 | Draft | *Oryza sativa* | Tracheophytes,<br>Angiosperms,<br>Monocots,<br>Commelinids,<br>Poales, Poaceae | Non-pathogenic | Mali: Africa | 2009 | 66.2 | 53 | 96.74/0.3 |
| *Xylella taiwanensis* PLS229<sup>T</sup> | 2.7 | 204<br>7 | Draft | Pear | Tracheophytes,<br>Angiosperms,<br>Eudicots, Rosids,<br>Rosales, Rosaceae | Pathogenic | Taiwan | - | 53 | 47 | 89.88/0.12 |
| *Xylella fastidiosa* ATCC 35879<sup>T</sup> | 2.5 | 210<br>5 | Draft | Grapevine | Tracheophytes,<br>Angiosperms,<br>Eudicots, Rosids,<br>Vitales, Vitaceae | Pathogenic | USA: Florida | 1987 | 51.8 | 49 | 99.23/0 |
| *Stenotrophomonas panacihumi* JCM 16536<sup>T</sup> | 3.9 | 349<br>7 | Draft | Soil of a ginseng field | - | Environmental | South Korea | 2010 | 68.8 | 51 | 97.59/0.34 |
| *Xanthomonas sontii* PPL1<sup>T</sup> | 4.8 | 393<br>7 | Draft | *Oryza sativa* | Tracheophytes,<br>Angiosperms, | Non-pathogenic | India | 2012 | 69 | 51 | 96.31/0.22 |
|  |  |  |  |  | Monocots,<br>Commelinids,<br>Poales, Poaceae |  |  |  |  |  |  |
| *Xanthomonas sacchari* CFBP 4641<sup>T</sup> | 4.9 | 409<br>8 | Draft | *Saccharum officinarum* | Tracheophytes,<br>Angiosperms,<br>Monocots,<br>Commelinids,<br>Poales, Poaceae | Pathogenic | Guadeloupe | 1980 | 69.1 | 51 | 100/0.41 |
| *Xanthomonas albilineans* CFBP 2523<sup>T</sup> | 3.6 | 296<br>7 | Draft | *Saccharum officinarum* | Tracheophytes,<br>Angiosperms,<br>Monocots,<br>Commelinids,<br>Poales, Poaceae | Pathogenic | Fiji | 1961 | 63.1 | 49 | 99.96/0.34 |
| *Xanthomonas hyacinthi* CFBP 1156 | 4.9 | 400<br>3 | Draft | *Hyacinthus orientalis* | Tracheophytes,<br>Angiosperms,<br>Monocots,<br>Asparagales,<br>Asparagaceae | Pathogenic | Netherlands | 1958 | 68.1 | 51 | 100/0.03 |
| *Xanthomonas translucens* DSM 18974<sup>T</sup> | 4.4 | 378<br>8 | Draft | *Hordeum vulgare* | Tracheophytes,<br>Angiosperms,<br>Monocots,<br>Commelinids,<br>Poales, Poaceae | Pathogenic | - | - | 67.7 | 53 | 98.5/0.12 |
| *Xanthomonas theicola* CFBP 4691 | 4.5 | 363<br>0 | Draft | *Camellia sinensis* | Tracheophytes,<br>Angiosperms,<br>Eudicots, Asterids,<br>Ericales, Theaceae | Pathogenic | Japan | 1974 | 68.4 | 49 | 99.95/0 |
| *Stenotrophomonas ginsengisoli* DSM 24757<sup>T</sup> | 3.3 | 300<br>1 | Draft | Soil from ginseng field | - | Environment<br>al | South Korea | 2010 | 65.9 | 76 | 98.72/0.76 |
| *Stenotrophomonas korensis* DSM 17805<sup>T</sup> | 3.0 | 269<br>0 | Draft | Compost | - | -- | South Korea | 2003 | 66.1 | 75 | 98.16/0.79 |
| *Stenotrophomonas terrae* DSM 18941<sup>T</sup> | 4.4 | 373<br>0 | Draft | Soil | - | Environment<br>al | Belgium | 2007 | 63.9 | 60 | 99.19/1.38 |
| *Stenotrophomonas humi* DSM 18929<sup>T</sup> | 4.1 | 359<br>1 | Draft | Soil | - | Environment<br>al | Belgium | 2007 | 64 | 62 | 99.6/1.03 |
| *Stenotrophomonas pictorum* JCM 9942<sup>T</sup> | 3.3 | 315<br>0 | Draft | Soil | - | Environment<br>al | - | 1928 | 66 | 61 | 99.84/0.32 |
| *Stenotrophomonas nitritireducens* DSM 12575<sup>T</sup> | 3.9 | 347<br>6 | Draft | Laboratory scale biofilters supplied with ammonia or dimethyl disulfide and ammonia | - | -- | Germany | 1997 | 68.3 | 54 | 95.5/0.78 |
| *Stenotrophomonas acidaminiphila* JCM 13310<sup>T</sup> | 3.9 | 351<br>1 | Draft | Sludge from a lab-scale anaerobic chemical waste water reactor | - | Environmental | Mexico | 1999 | 68.8 | 56 | 96.69/1.51 |
| *Stenotrophomonas daejeonensis* JCM 16244<sup>T</sup> | 3.2 | 288<br>8 | Draft | Sewage water | - | Environmental | South Korea | 2010 | 68.6 | 56 | 99.59/0.86 |
| *stenotrophomonas hibiscicola* ATCC 19867<sup>T</sup> | 4.4 | 392<br>8 | Draft | Plant | - | - | - | - | 66.4 | 64 | 100/0 |
| *Stenotrophomonas pavanii* DSM 25135<sup>T</sup> | 4.3 | 378<br>3 | Draft | Stems of sugar cane | Tracheophytes, Angiosperms, Monocots, Commelinids, Poales, Poaceae | Symbiotic | Brazil | 2011 | 66.2 | 67 | 100/0 |
| *Stenotrophomonas africana* LMG 22072<sup>T</sup> | 4.5 | 399<br>1 | Draft | Cerebrospinal fluid | - | Opportunistic pathogen | Democratic Republic of the Congo | 1994 | 66.3 | 65 | 97.05/0.23 |
| *Pseudomonas geniculate* JCM 13324<sup>T</sup> | 4.8 | 433<br>9 | Draft | Tap water | - | Environmental | USA | 1895 | 66.2 | 66 | 99.76/0.41 |
| *Stenotrophomonas maltophilia* NCTC 10257<sup>T</sup> | 5.0 | 449<br>1 | Complete | Mouth | - | Opportunistic pathogen | Germany | 1900/1961 | 66.1 | 73 | 100/0.34 |
| *Stenotrophomonas beteli* LMG 00978<sup>T</sup> | 4.4 | 390<br>7 | Draft | *Piper betle* | Tracheophytes, Angiosperms, Magnoliids, Piperales, Piperaceae | - | Sri Lanka | 1928 | 66.8 | 67 | 99.08/0 |
| *Stenotrophomonas indicatrix* WS40<sup>T</sup> | 4.5 | 406<br>7 | Draft | Dirty dishes | - | Environmental | Germany: Bonn | 2013 | 66.4 | 65 | 99.91/0.57 |
| *Stenotrophomonas lactitubi* M15<sup>T</sup> | 4.8 | 437<br>8 | Draft | Milking machine biofilm | - | - | Germany: Koenigswinter-Vinzel | 2014 | 65.9 | 66 | 100/0.41 |
| *Stenotrophomonas chelatiphaga* DSM 21508<sup>T</sup> | 3.9 | 342<br>0 | Draft | Municipal sewage sludge | - | Environmental | Russia | 2009 | 66.9 | 62 | 98.75/0.04 |
| *Stenotrophomonas bentonitica* VV6 | 4.3 | 3805 | Draft | Arctic sea water near B02 station | - | Environmental | - | - | 66.5 | 60 | 99.91/1.93 |
| *Stenotrophomonas rhizophila* DSM 14405<sup>T</sup> | 4.6 | 3971 | Draft | Root | - | Symbiotic | Germany: Rostock | - | 67.3 | 67 | 100/0.21 |
| *Pseudoxanthomonas composti* GSS 15<sup>T</sup> | 4.3 | 3664 | Draft | Compost | - | Environmental | China | - | 68.2 | 52 | 99.31/0.8 |
| *Pseudoxanthomonas spadix* DSM 18855<sup>T</sup> | 3.3 | 3042 | Draft | Oil-contaminated soil | - | Environmental | Taiwan | Before 2006 | 67.8 | 48 | 98.97/0.59 |
| *Pseudoxanthomonas daejeonensis* DSM 17801<sup>T</sup> | 3.5 | 3143 | Draft | Soil from a ginseng field | - | Environmental | - | - | 68.8 | 58 | 99.66/0.11 |
| *Pseudoxanthomonas kaohsiungensis* DSM 17583<sup>T</sup> | 3.7 | 3420 | Draft | Oil-polluted site | - | Environmental | Taiwan | 2003 | 69.7 | 52 | 99.66/1.42 |
| *Pseudoxanthomonas koreensis* KCTC 12208<sup>T</sup> | 3.0 | 2681 | Draft | Soil from a ginseng field | - | Environmental | South Korea | - | 70.1 | 52 | 99.48/0.04 |
| *Pseudoxanthomonas jiangsuensis* DSM 22398<sup>T</sup> | 3.7 | 3389 | Draft | DDT-contaminated soil | - | Environmental | China | 2008 | 70.3 | 50 | 99.31/1.03 |
| *Pseudoxanthomonas broegbernensis* DSM 12573<sup>T</sup> | 3.5 | 3024 | Draft | Ammonia-supplied biofilters | - | Environmental | Germany | Before 1998 | 70.6 | 54 | 99.66/1.74 |
| *Pseudoxanthomonas suwonensis* DSM 17175<sup>T</sup> | 3.4 | 3071 | Draft | Cotton waste composts | - | Environmental | South Korea | Before 2005 | 70.3 | 55 | 99.66/0.34 |
| *Pseudoxanthomonas taiwanensis* DSM 22914<sup>T</sup> | 3.0 | 2729 | Draft | Hot spring | - | Environmental | Taiwan | Before 2009 | 72.08 | 56 | 99.16/0.34 |
| *Pseudoxanthomonas sangjuensis* DSM 28345<sup>T</sup> | 3.2 | 2908 | Draft | Greenhouse soil | - | Environmental | Republic of Korea | Before 2014 | 68.6 | 51 | 99.95/0.46 |
| *Pseudoxanthomonas kalamensis* DSM 18571<sup>T</sup> | 3.0 | 2697 | Draft | Soil contaminated with polycyclic aromatic hydrocarbons & polychlorinated biphenyls | - | Environmental | United States of America | Before 2006 | 65.9 | 48 | 99.84/1.5 |
| *Pseudoxanthomonas dokdonensis* DSM 21858<sup>T</sup> | 3.5 | 3153 | Draft | Soil | - | Environmental | South Korea | Before 2008 | 64.4 | 50 | 99.59/0.41 |
| *Pseudoxanthomonas wuyuanensis* DSM 100640<sup>T</sup> | 4.6 | 4003 | Draft | Saline-alkaline soil | - | Environmental | China | Before 2015 | 65.7 | 50 | 100/1.08 |
| *Pseudoxanthomonas mexicana* DSM 17121<sup>T</sup> | 3.9 | 3660 | Draft | Anaerobic digester | - | Environmental | Mexico | Before 2005 | 67.4 | 54 | 99.49/1.71 |
| *Pseudoxanthomonas arseniciresistens* KAS 5_3<sup>T</sup> | 3.9 | 4049 | Draft | Arsenic contaminated ground water | - | Environmental | India | - | 66.5 | 52 | 99.95/1.19 |
| *Pseudoxanthomonas japonensis* DSM 17109<sup>T</sup> | 4.0 | 3673 | draft | Urban riverside soil |  | Environmental | Japan | Before 2005 | 67.3 | 50 | 99.95/0.76 |
| *Pseudoxanthomonas indica* P15<sup>T</sup> | 3.9 | 3593 | Draft | Open hexachlorocyclohexane dumpsite soil | - | Environmental | India | - | 65.4 | 49 | 99.89/0.47 |
| *Pseudoxanthomonas yeongjuensis* DSM 18204<sup>T</sup> | 3.9 | 3389 | Draft | Soil of a ginseng field | - | Environmental | Republic of Korea | 2005 | 65.1 | 53 | 99.95/0.76 |
| *Pseudoxanthomonas gei* KCTC 32298<sup>T</sup> | 3.4 | 3122 | Draft | Stem of *Geum aleppicum* Jacq | Tracheophytes, Angiosperms, Eudicots, Rosids, Rosales, Rosaceae | Endophyte | Taibai Mountain in Shaanxi Province, north-west China | - | 65.4 | 45 | 99.95/0.41 |
| *Pseudoxanthomonas sacheonensis* DSM 19373<sup>T</sup> | 4.0 | 3582 | Draft | BTEX-contaminated soil | - | Environmental | Republic of Korea | Before 2007 | 64.3 | 50 | 100/0.91 |

### Phylogenomics of XSXP phylogroup

To access the relationship amongst the strains of XSXP phylogroup, we constructed and compared genome based phylogeny at the two levels. Phylogeny constructed on the basis of large set of genes core to the bacterial world (figure 1) and set of genes core to XSXP phylogroup (figure 2) correlated with each other (pl. see methods). Based on our previous study of order *Xanthomonadaceae* (*Lysobacteraceae*), we used *Luteimonas mephitis* DSM12574^T^ as an outgroup of XSXP (Kumar, Bansal et al. 2019). Here, amongst the XSXP phylogroup, *Pseudoxanthomonas* members were more diverse and ancestral to other three genera. Interestingly, other three genera formed two mega species groups (MSG) i.e. plant pathogens *Xanthomonas* and *Xylella* comprised one MSG referred as XX-MSG and *Stenotrophomonas* formed another MSG referred as S-MSG. This analysis also revealed that both the MSGs comprised of species complexes. XX-MSG consist of three species complexes, *X. citri* complex (Xcc), *X. hyacinthi* complex (Xhc) and *Xylella fastidosa* complex (Xfc). Xcc comprises of at least 27 species that are primarily pathogens of dicot plants. Xhc comprises of at least 6 species that are primarily pathogen of monocot plants. Interestingly, *Stenotrophomonas panacihumi*, an environmental species, reflected as singleton phylogenetic outlier of Xcc. *Xylella* as a species complex or minor variant lineage sandwiched in between both of the *Xanthomonas* complexes i.e. Xcc and Xhc. In case of S-MSG, we found one species complex i.e. *Stenotrophomonas maltophilia* complex (Smc) comprising of two nosocomial species i.e. *S. maltophilia* and *S. africana.*

**Figure 1:**
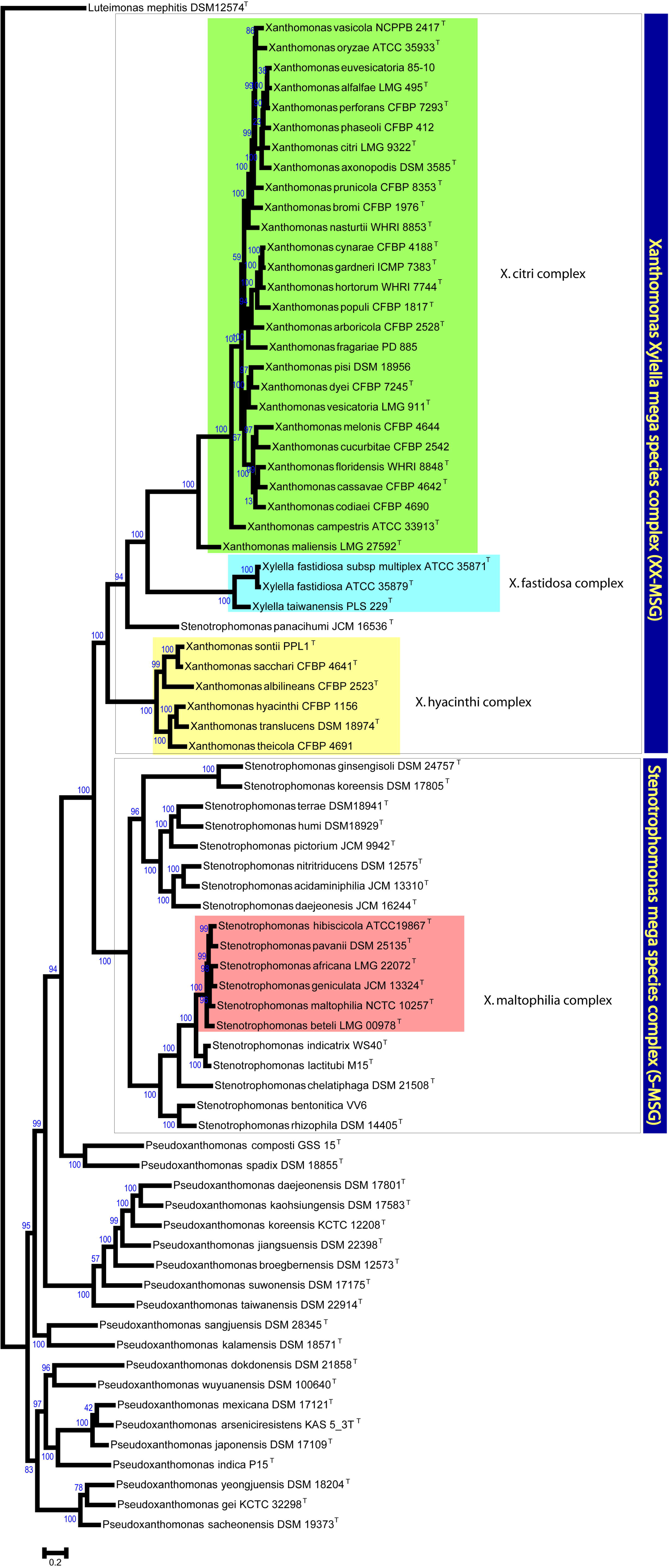
Phylogenetic tree obtained using PhyloPhlAn for XSXP phylogroup. Both the mega species groups are represented by blue boxes and four species complexes are represented by colored boxes. *Luteimonas mephitis* DSM12574^T^ was used as an outgroup and bootstrap values are mentioned on the nodes.

**Figure 2:**
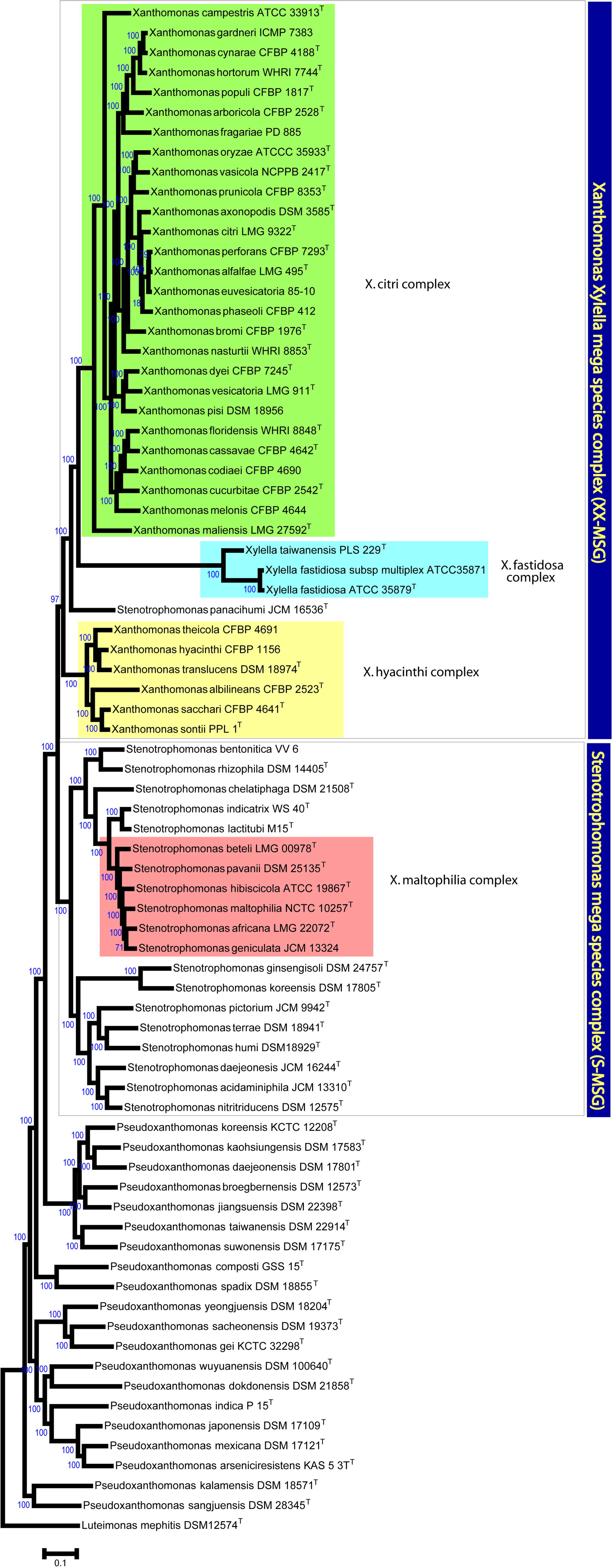
Phylogenetic tree based on core genome phylogeny of XSXP phylogroup. Both the mega species groups are represented by blue boxes and four species complexes are represented by colored boxes. *Luteimonas mephitis* DSM12574^T^ was used as an outgroup and bootstrap vales are mentioned on the nodes.

### Taxonogenomics of XSXP phylogroup

In order to delineate members of XSXP at genera and species complexes level, we further carried out taxonogenomic analysis using AAI (figure 3), POCP (figure 4) and at the level of species using ANI (supplementary figure 1). Due to drastic genome reduction in case of *Xylella* resulting in reduction of coding sequences, POCP is not suitable for *Xylella* (Hayashi Sant’Anna, Bach et al. 2019). Interestingly, according to the genus cut-off value of AAI all the strains belong to the same genus i.e., *Xanthomonas* and hence need to name as *Xanthomonas.* Similarly, according to POCP cut-off all members of *Xanthomonas, Stenotrophomonas* and *Pseudoxanthomonas* belong to one genus. Furthermore, within XX-MSG, 27 species formed Xcc while other 6 species formed Xhc. Third complex that is intermediate to Xcc and Xhc is a variant *Xanthomonas (Xylella) fastidosa* complex (Xfc) consisting by 2 species of *Xylella.* Here, members of Xcc are pathogens of dicots, except 5 species which have monocot hosts i.e. *X. vasicola, X. oryzae, X. axonopodis, X. bromi* and *X. maliensis*. On the other hand, members of Xhc are pathogens or associated with monocots, except *X. theicola* infecting dicots. In case of S-MSG, the *Xanthomonas (Stenotrophomonas) maltophilia* complex (Xmc) consisting of 6 species with two of them being clinical origin i.e. *S. maltophilia* and *S. africana*. Further, ANI values clearly depicted that all the members belong to different species except for *X. alfalfa, X. perforans, X. euvesicatoria* and *X. gardneri, X. cynarae*, which represent miss-classified different species.

**Figure 3:**
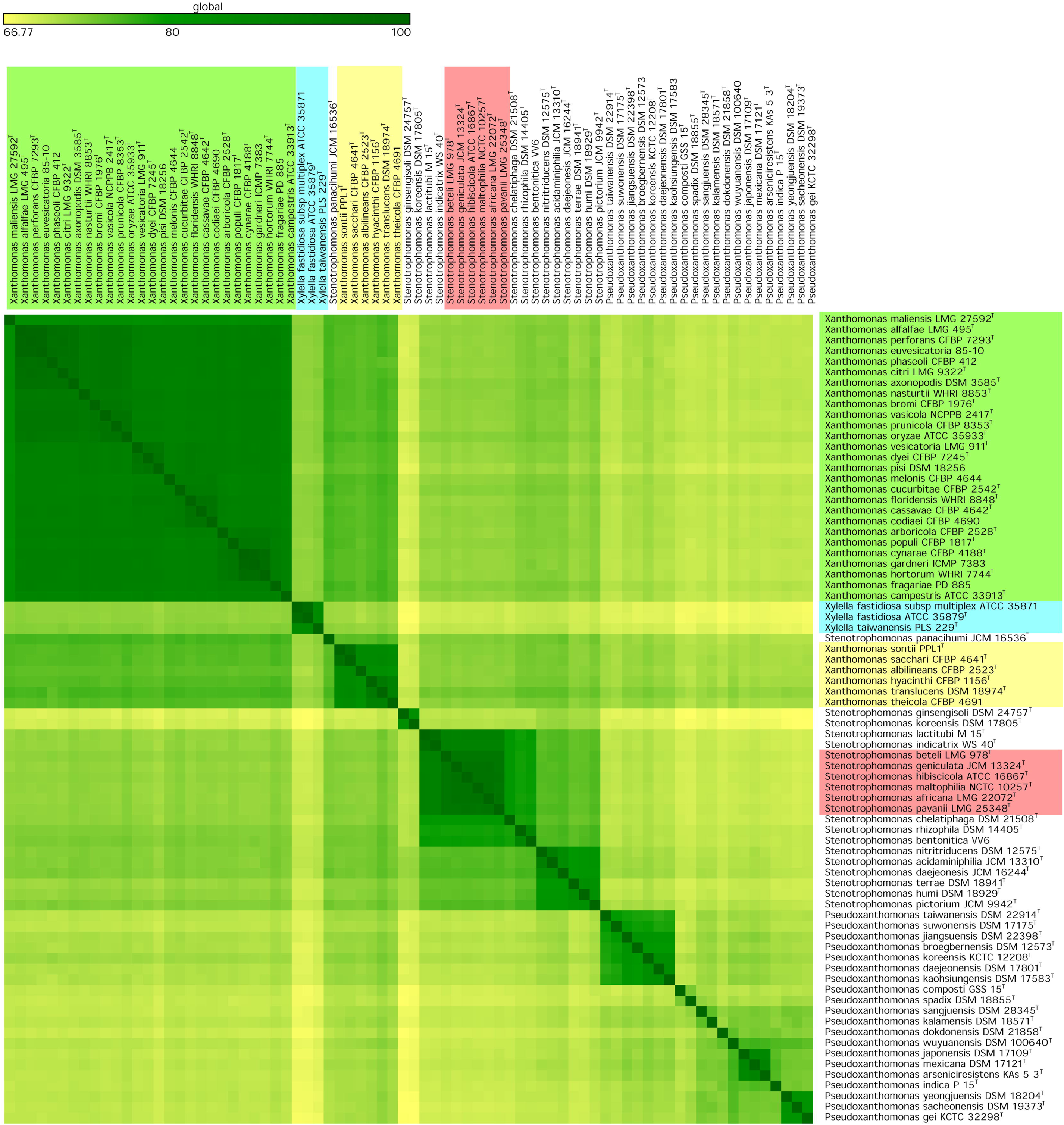
Heatmap of genome similarity matrix showing average amino acid identity (AAI) Values. All the species complexes are represented in the colored boxes.

**Figure 4:**
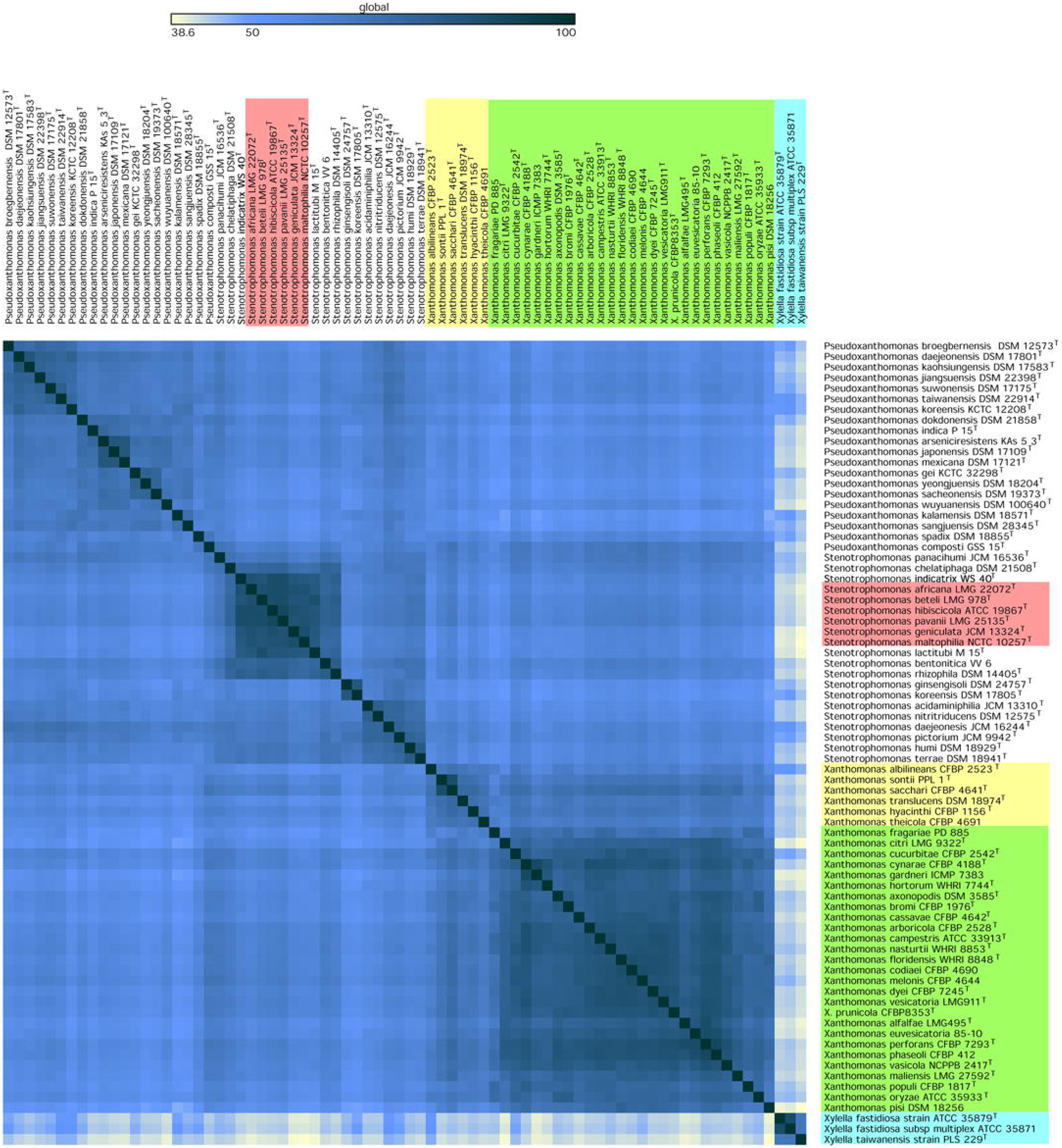
Heatmap showing percentage of conserved proteins (POCP). All the species complexes are represented in the colored boxes.

### Xanthomonadin and exopolysaccharide cluster

Distinct yellow xanthomonadin pigment, encoded by *pig* gene cluster, and thick mucous exopolysaccharide or xanthan encoded by *gum* gene cluster are characteristic features of canonical plant associated *Xanthomonas* species (Rajagopal, Sundari et al. 1997, Katzen, Ferreiro et al. 1998). Since, we are expanding breadth of *Xanthomonas* genus on the basis of phylo-taxono-genomics parameters, we scanned these clusters in the genomes of XSXP constituent genera and species (figure 5). Among the XX-MSG, both the clusters were highly conserved in sequence and distribution in Xcc and Xhc. Xhc members have diversified clusters with *X. albilineans* and *X. theiocola* as exceptions as they lack *gum* gene cluster. Further, members of Xfc showed incomplete and possibly degenerated clusters, with *gumN, gumM, gumI, gumG* and *gmuF* absent from the *gum* cluster and orfs 8, 11, 13 and 14 absent from the pigment cluster.

**Figure 5:**
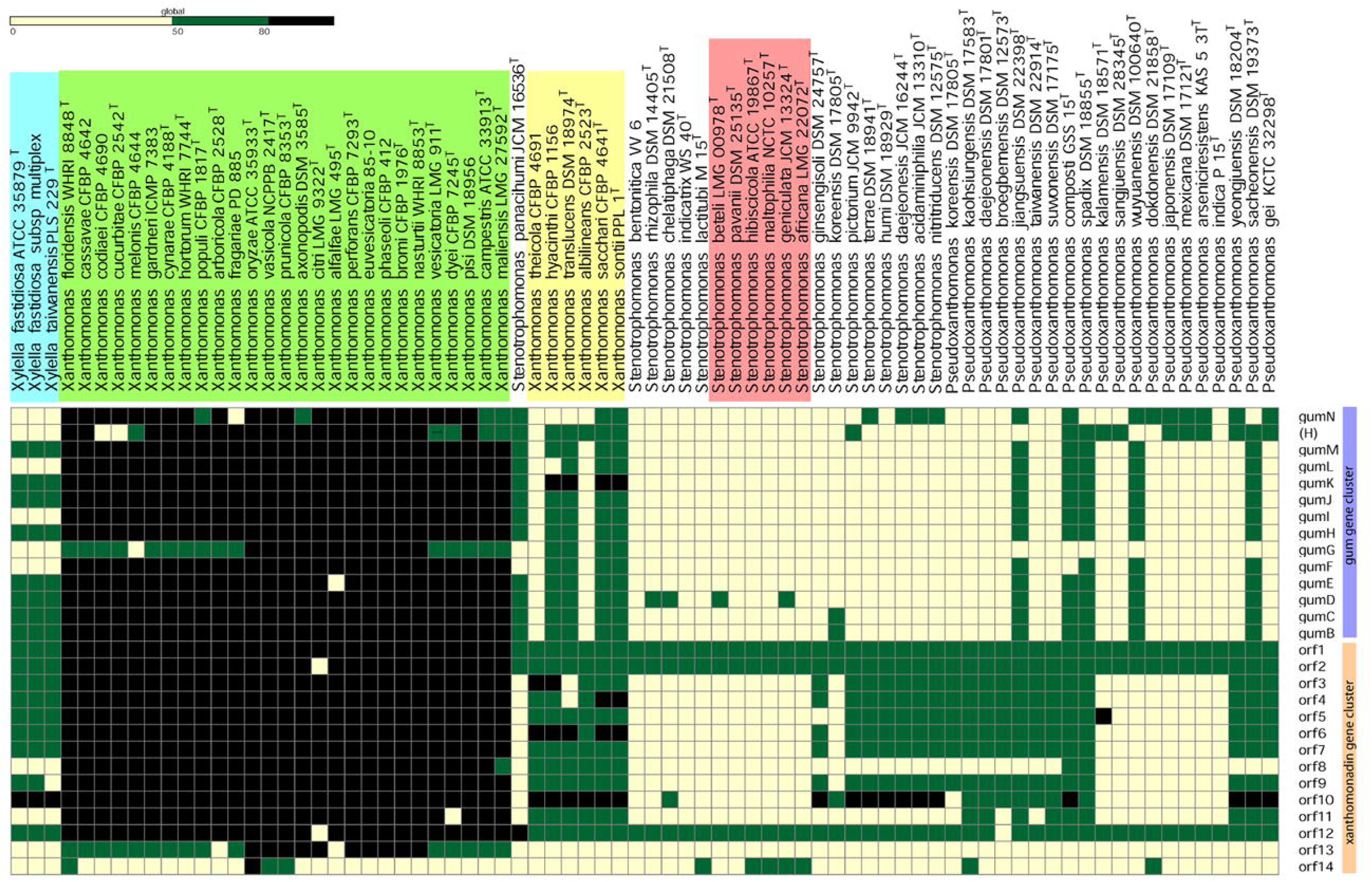
Heatmap showing status of xanthomonadin pigment and *gum* cluster in XSXP phylogroup.

**Figure 6:**
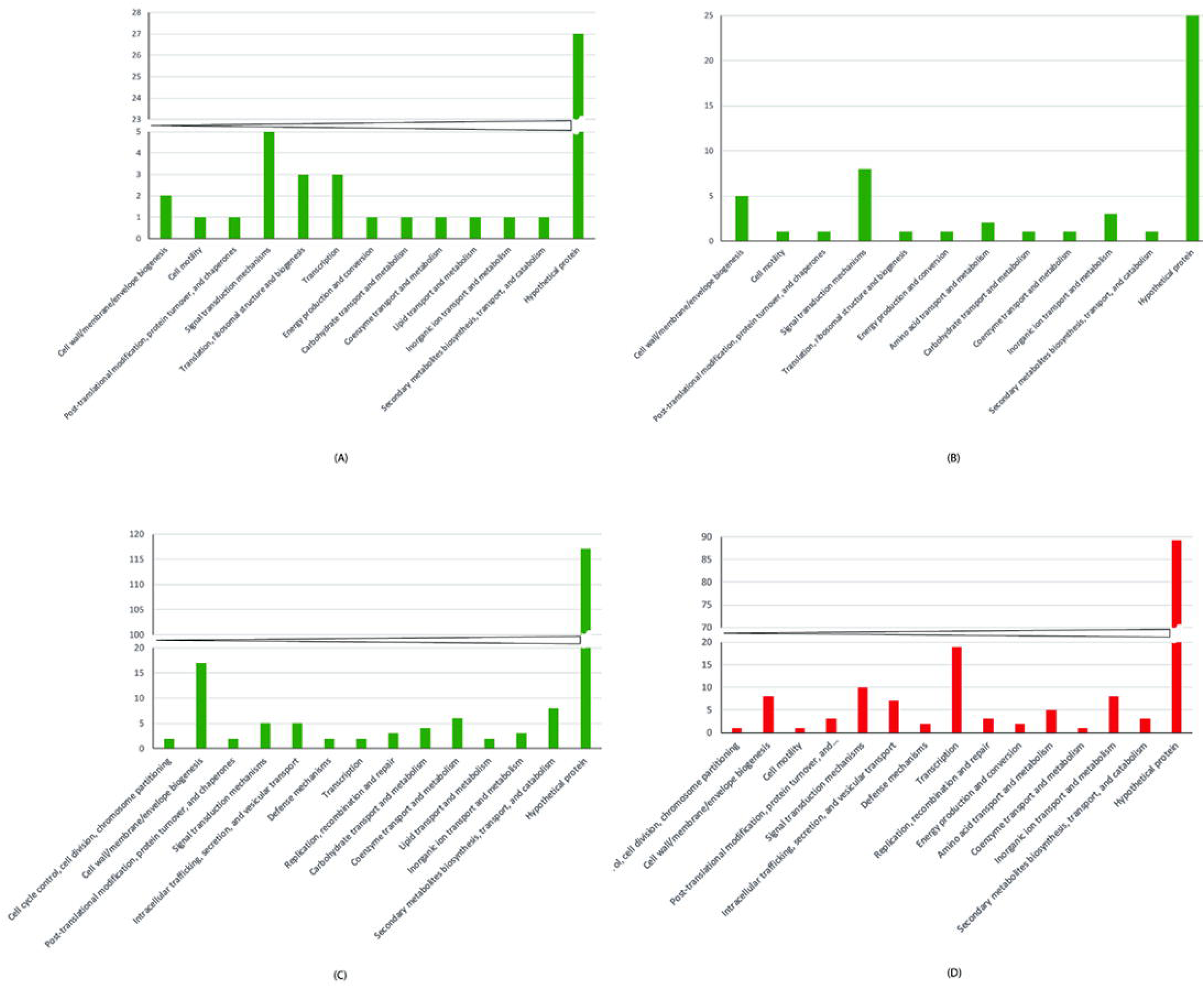
Distribution of COG-based functional categories of unique genes. (A) *X. citri* complex, (B) *X. hyacinthi* complex, (C) *X. fastidosa* complex and (D) *S. maltophilia* complex.

Interestingly, in large majority members of other two genera understudy, these clusters are either absent or incomplete and less conserved. Specifically, while all members of genus *Stenotrophomonas* lacks *gum* gene cluster, and 5 out of 20 species of *Pseudoxanthomonas* i.e. *P. jiangsuensis, P. composti, P. spadix, P. wuyuanensis* and *P. sacheonensis* harbour *gum* gene cluster. Whereas, xanthomonadin cluster is widely present in other two genera i.e. 7 out of 19 *Stenotrophomonas* and 13 out of 20 *Pseudoxanthomonas* species (figure 5). Interestingly, none of the Xmc species have both the clusters.

### Pan genome analysis

Since XSXP group comprises of species complexes of diverse ecological niches, we performed pan genome analysis to access the species complex specific gene content. Overall, 47,487 pan genes and 399 core genes were detected for the XSXP phylogroup. Further, 48, 53, 205 and 175 genes were unique to Xcc, Xhc, Xfc and Xmc respectively. Inspection of Xmc unique genes revealed a unique type II secretion system, peroxidase, peptidases, efflux pumps and transporters like antibiotics/antimicrobials, fluoride ions, solvent, TonB dependent receptors, transcriptional regulators etc. (supplementary table 1). Further, unique genes of Xfc belong to adhesin like type IV pili formation apart from glycosyltransferases, methyltransferases etc. (supplementary table 2). Overall COG classification revealed that hypothetical proteins are dominant class in Xcc, Xhc, Xfc and Xmc suggesting unknown functions playing role in their success. Interestingly, in both Xcc and Xhc, COG class related to “signal transduction mechanisms” is second major class after hypothetical proteins, Xfc is having more of “cell wall/membrane/envelope biogenesis”, “secondary metabolites biosynthesis, transport and catabolism” and “coenzyme transport and metabolism”. Whereas, Xmc was having more of “transcription”, “signal transduction mechanisms”, “intracellular trafficking, secretion and vesicular transport”, “cell wall/membrane/envelope biogenesis” and “inorganic ion transport and metabolism”. However, “intracellular trafficking, secretion, and vesicular transport”, “defence mechanism”, “replication, recombination and repair” and “cell cycle control, cell division, chromosome partitioning” were present in both Xmc and Xfc.

## Discussion

The ability to sequence genome of large number of strains in a cost effective manner is transforming the way we do genetics, phylogeny and taxonomy of bacteria. Inferring phylogeny based on limited sequence information like 16S rRNA and housekeeping genes can be highly misleading (Sangal, Goodfellow et al. 2016). Genomic investigation is the finest and best way to look at organism identity, biology and ecology in proper context. Genome based comprehensive taxonomic studies at various levels such as: order *Methylococcales*, order *Bacillales*, genus *Borrelia*, genus *Lactobacillus* have resulted in resolving chaos and provided basis for reclassification (Orata, Meier-Kolthoff et al. 2018, Salvetti, Harris et al. 2018, De Maayer, Aliyu et al. 2019, Margos, Fingerle et al. 2019). Similarly, in whole genome studies using representative reference strains from our lab have successfully resolved misclassifications in order *Xanthomonadales* (*Lysobacterales*) at the level of family (Kumar, Bansal et al. 2019). Similarly, in case of plant pathogenic *Xanthomonas* population at the level of species and clones (Bansal, Midha et al. 2017, Kumar, Bansal et al. 2019), while in case of *Stenotrophomonas* at the level of species (Patil, Kumar et al. 2018). However, deeper genera and intra-genera genome based investigation at phylogenetic and taxonomic level was lacking. Considering importance of members of XSXP phylogroup in plant and human health apart from biotechnological potential, we carried out in-depth investigation by incorporating large or all the members of four closely related genera through modern genome-based criteria.

Present taxonogenomic and phylogenomic analysis revealed the role of evolutionary and ecological diversity in formation of species and species complexes that actually belong to genus *Xanthomonas* but mistakenly represented into distinct genera as *Xylella, Stenotrophomonas* and *Pseudoxanthomonas*. The basal and multiple groups constituting species of *Pseudoxanthomonas* suggests it to be ancestral to other three genera. Hence, it is not surprising that this large genus is named as *Pseudoxanthomonas* (Finkmann, Altendorf et al. 2000). At the same time, members of other three genera form phylogenetic mega species groups constituting species complexes from diverse ecological niches indicates much recent origin and ongoing specialization. This is quite obvious in canonical plant associated and pathogenic *Xanthomonas* species that form a mega-group distinct from *Stenotrophomonas* and *Pseudoxanthomonas* whose members are highly versatile. This suggests the role of lifestyle in divergence and evolution into distinct groups that got reflected as separate taxonomic units at the level of genera/family using limited genotypic and phenotypic data. Most importantly, apart from mega species groups in XSXPs, the clarity and robustness was clear in understanding relationship among XSXP suggesting importance of deep phylogenetic studies across genera. In our case this was possible because of the phylogenetic foundation provided by species of *Pseudoxanthomonas* for other three genera, indicating importance of genomic resource of *Pseudoxanthomonas.*

Establishing position of *Xylella* with fastidious nature and extremely reduced GC content is of critical importance. While finding that at taxonogenomic level, all the four genera are not distinct but belong to one genus, the element of surprise is that genus *Xylella* is more related to *Xanthomonas* than *Stenotrophomonas*. It revealed as a variant lineage or species complex of *Xanthomonas*, which was further confirmed by the sandwiched phylogenomic position of the *Xylella* within the XX-MSG. Overall, XX-MSG seems to have evolved into three species complexes. One corresponding to Xhc which consists of either non-pathogenic species *X. sontii* or other that infect monocots with *X. theicola* as an exception. Second complex corresponds to *X. citri* complex Xcc that primarily consists of pathogenic species infecting dicots with few exceptions and third is Xfc, a phytopathogen of fastidious nature. Also, while XX-MSG is primarily plant associated or plant pathogenic, the S-MSG consist of members that have diverse lifestyle as soil dwelling, aquatic, plant epiphyte and even as nosocomial but not pathogenic to plants (Patil, Midha et al. 2016). Clinical strains of *S. maltophilia* were found to belong to species complex (Patil, Kumar et al. 2018, Gröschel, Meehan et al. 2019). Accordingly, we can find a species complex in S-MSG consisting of clinical species along with plant and environmental associated *Stenotrophomonas* species. However, this need to rename into *Xanthomonas maltophilia* complex (Xmc) in the light of new findings from this study.

Conservation of xanthomonadin pigment and *gum* gene clusters in Xcc and Xfc of the XX-MSG suggests importance of pigment and exopolysaccharide production in obligate plant associated lifestyle. Since, plants are directly and continuously exposed to light, for a successful phytopathogen, it is important to have a unique pigment like xanthomonadin (Rajagopal, Sundari et al. 1997). It is shown that xanthomonadin provides protection against photodamage in *X. oryzae* pv. oryzae that infect rice (Rajagopal, Sundari et al. 1997). Since plants are also known to produce huge amount of antimicrobial compounds and secondary metabolites (Ramírez-Gómez, Jiménez-García et al. 2019), there is need to have unique and thick polysaccharide coat. In fact, highly mucoid and distinct yellow coloured colonies reflect and supports our hypothesis that these clusters are highly evolved in canonical *Xanthomonas* species from XX-MSG. Interestingly, both these clusters were found to be diversified in Xhc as compared to Xcc, this can be explained by majorly dicot associated lifestyle of Xcc as compared to majorly monocot associated lifestyle of Xhc. The leaves of monocot and dicots are different which may affect the penetration effects on bacteria, though this hypothesis needs further studies. Importantly, both the clusters were degenerated or incomplete in case of Xfc and that can be supported by dual lifestyle of *Xylella* that occurs within insect and plant where it is directly injected into xylem (Chatterjee, Almeida et al. 2008). Since, *Xylella* is never exposed to light and plant defence response unlike *Xanthomonas* (i.e. Xcc and Xhc) it is not surprising that in *Xylella* the clusters are not under natural selection. Genetic studies revealed that *pig* cluster is not required for virulence in plants (Rajagopal, Sundari et al. 1997). Hence, conservation of *pig* cluster in Xcc and Xhc without their role in virulence suggests that epiphytic mode is also important in plant adaptation (Pandey and Sonti 2010).

Interestingly, presence of a *pig* and *gum* clusters in few members of XSXP outside XX-MSG suggests that these clusters are ancestral and present as incomplete clusters in the population even before emergence of Xcm and Xch. Later on, they got evolved and selected when members of XSXP or ancestor of XX-MSG came in contact with plants. The ancestor of XX-MSG had probably both the clusters and advantage for plant-associated lifestyle. Obligate plant associated lifestyle of mega complex of *Xanthomonas* suggests that selection correlates with sudden emergence and radiation of angiosperms. At the same time, presence of a primitive, ancestral and incomplete pigment and *gum* gene clusters suggest that these clusters are not of foreign origin but existed in ancestral XSXP population. However, with the emergence of plant kingdom and mysterious radiation of angiosperms can find correlation with emergence of plant associated XX-MSG (Friedman 2009). Presence of monocots and dicots associated *Xanthomonas* complexes suggests further selection as and when monocots and dicots arrived in the scene. Importance of pigment cluster in case of *Xanthomonas* phytopathogens can be compared with pivotal role of chlorophyll pigment in emergence of plants. Hence, both in plants and their major bacterial pathogen, unique pigments have played role in their emergence. While in plants the pigment is to harvest light for photosynthesis, but in pathogens, a pigment is for protection against photodamage.

While Darwinian selection acting on minor or random mutations is well established since classical experiment of Luria–Delbrück experiment (Jones, Thomas et al. 1994), our XSXP analysis provides evidence on how evolution acts on existing variations that too to large scale variation. Horizontal gene transfers are dominant force that drive bacteria evolution and there are numerous evidences (De la Cruz and Davies 2000, Ochman, Lawrence et al. 2000). Such cases may lead to more of opportunistic pathogens and in short-time scale. However, in XSXP story we see selection happening on existing on large-scale variation and further systematic selection that led to probably one of the highly successful plant associated or plant pathogenic group of bacteria with specificity at the level of host and tissues. Our DEEPT genomics not only reveal established MSGs, particularly co-evolved XX-MSG but also reveal variant or opportunistic lineages or species complexes i.e. Xfc and Xmc mediated through more recent horizontal gene transfers as seen by relatively large number of unique genes. This indicates importance of acquisition of genes for fastidious nature of Xfc and opportunistic pathogenic lifestyle of clinical pathogens of Xmc.

Interestingly, even though highly reduced genome compared to other XSXP members, *Xylella* have acquired more genes than Xmc. Even though *Xylella* seems to have underwent drastic genome reduction, acquisition of large number of unique genes of particular functions has played important role in its emergence into a successful phytopathogen. In both these variant lineages or complexes (Xfc and Xmc), acquisition of functions related to regulation suggest that apart from gene gain, the regulation of core or novel genes has also played important role in their success. Commonality of functions of unique genes related to functions like “intracellular trafficking, secretion, and vesicular transport”, “defence mechanism”, “replication, recombination and repair” and “cell cycle control, cell division, chromosome partitioning” again reiterate their opportunistic, variant origin and parallels in evolution for dual lifestyle. Even though both have dual life style, unique gene analysis allowed us to pinpoint functions like adhesion in *Xylella* and efflux proteins/peroxidise, etc. (for antimicrobial resistance, particularly in hospital settings) along with a novel type II secretion system in Xmc that were important in their success. Even though an opportunistic human pathogen, a novel type II secretion system may be compensating the absence of type III secretion system in Smc.

Even within *Stenotrophomonas* group or S-MSG, formation of a species complex corresponding to MDR nosocomial pathogenic suggest ongoing evolution and diversification as seen in case of *Xylella* within XX-MSG. This finding is valuable in further understanding this emerging opportunistic human pathogen. In this context, the clinical pathogenic species that form complex and referred as *Stenotrophomonas maltophilia* complex or Smc in earlier studies can be referred to as *Xanthomonas maltophilia* complex or Xmc. At the same time unique genes in Xcc and Xhc may be related to adaptation to dicots and monocots respectively. While large number of unique genes encode hypothetical protein suggesting importance of further functional genetic studies in Xcc, Xhc, Xfc and Xc, other major classes also provide clue regarding their evolution. For example, importance of signal transduction in Xcc and Xhc, which then can important targets for both basic and applied studies. More systematic gene content analysis by excluding variants within this group will also allow us to obtain insights into genes important for adaptation to monocots and dicots. Overall, our study not only reiterates power and accuracy of systematic and large scale or deep taxonogenomics but also has major implications in how we manage and study the members of any genera of high importance in agriculture, medicine and industry. Such deep phylo-taxono-genomics approaches or what we like to refer as DEEPT (as depth) genomic studies in understanding, exploiting and managing good/bad bacteria by zeroing on stable, unique and novel (STUN) markers. Hence, such a broader phylo-taxono-genomic studies can be deeply stunning (DEEPT-STUN).

### Emended Description of Genus *Xanthomonas* (Dowson 1939)

Synonym: *Pseudoxanthomonas* (Finkmann, Altendorf et al. 2000), *Stenotrophomonas* (Palleroni and Bradbury 1993) and *Xylella* (Wells, Raju et al. 1987).

Based on whole genome based taxonomic criteria like AAI and POCP along with phylogenetic analysis with set of genes conserved in bacterial world and genes core to members of four genera i.e. *Xanthomonas, Xylella, Stenotrophomonas* and *Pseudoxanthomonas.* The genus *Xanthomonas* comprise of all the reported species of *Xylella, Stenotrophomonas* and *Pseudoxanthomonas.* Members of the genus are from diverse ecological niches and follow diverse lifestyle. The GC content varies from 52% to 76 % and genome size varies from 2.5 Mb to 5.5 Mb. This comprises of at least four species complexes i.e. *X. citri* complex, *X. hyacinthi* complex, *X. fastidosa* complex and *X. maltophilia* complex with distinct features as mentioned below.

The type species is *Xanthomonas campestris*.

### Emended description of *Xanthomonas citri* complex

This complex was identified based on whole genome taxonomic criteria like ANI, AAI and POCP analysis along with phylogenetic analysis with set of genes conserved in bacterial world and genes core to members of four genera i.e. *Xanthomonas, Xylella, Stenotrophomonas* and *Pseudoxanthomonas.* The complex comprises 27 species namely, *X. vasicola, X. oryzae, X. euvesicatoria, X. alfalfa, X. perforans, X. phaseoli, X. citri, X. axonopodis, X. prunicola, X. bromi, X. nasturtii, X. cynarae, X. gardneri, X. hortorum, X. populi, X. arboricola, X. fragariae, X. pisi, X. dyei, X. vesicatoria, X. melonis, X. cucurbitae, X. floridensis, X. cassava, X. codiaei, X. campestris, X. maliensis.* All of these are devastating plant pathogens, primarily of dicots with exception of *X. oryzae, X, vasicola, X. bromi, X. maliensis* and *X. axonopodis* which have monocot hosts as described in Swings et al. (Van den Mooter and Swings 1990).

### Emended description of *Xanthomonas hyacinthi* complex (Xhc)

This complex was identified based on whole genome taxonomic criteria like ANI, AAI and POCP analysis along with phylogenetic analysis with set of genes conserved in bacterial world and genes core to members of four genera i.e. *Xanthomonas, Xylella, Stenotrophomonas* and *Pseudoxanthomonas.* The complex comprises 6 species namely, *X. hyacinthi, X. sontii, X. sacchari, X. albilineans, X. translucens* and *X. theicola.* These are primarily associated with monocots following pathogenic and non-pathogenic lifestyle. However, *X. theicola* is an exception as it is pathogenic to dicot hosts. The description is based on Vauterin et al. (Vauterin, Hoste et al. 1995).

### Emended description of *Xanthomonas fastidosa* complex (Xfc)

Synonym: *Xylella fastidosa*

This complex was identified based on whole genome taxonomic criteria like ANI, AAI and POCP analysis along with phylogenetic analysis with set of genes conserved in bacterial world and genes core to members of four genera i.e. *Xanthomonas, Xylella, Stenotrophomonas* and *Pseudoxanthomonas.* The complex comprises 2 species namely, *Xylella fastidosa* and *Xylella taiwanensis* which are plant pathogens. According to whole genome information, average GC content is 52.4 % and average genome size is 2.6 Mb. The description is based on Wells et al. (Wells, Raju et al. 1987).

### Emended description of *Xanthomonas maltophilia* complex (Xmc)

Synonyms: *Stenotrophomonas maltophilia* complex (Smc)

This complex was identified based on whole genome taxonomic criteria like ANI, AAI and POCP analysis along with phylogenetic analysis with set of genes conserved in bacterial world and genes core to members of four genera i.e. *Xanthomonas, Xylella, Stenotrophomonas* and *Pseudoxanthomonas.* The complex comprises 6 species namely, *S. hibiscicola, S. pavanii, S. africana, S. geniculata, S. maltophilia* and *S. beteli.* Out of these, only *S. maltophilia* and *S. africana* are of nosocomial origin. The description of this complex is same as described by Palleroni and Bradbury (Palleroni and Bradbury 1993).

## Methods

### Bacterial strains and culture conditions

A total of 76 strains were used in the present study. Genome sequence of 33, 19 and 2 were type strains of *Xanthomonas, Stenotrophomonas* and *Xylella*, were available from the NCBI database (https://www.ncbi.nlm.nih.gov/). Out of twenty type strains of *Pseudoxanthomonas* we have procured and generated whole genome information of fifteen strains as genomes of remaining five type strains was publically available (https://www.ncbi.nlm.nih.gov/). The type strains were procured from culture collections of The Leibniz Institute DSMZ – German Collection of Microorganisms and Cell Cultures GmbH DSMZ and Korean Collection for Type Cultures (KCTC, Korea) KCTC. All the strains were grown on the media and under conditions recommended by respective culture collections.

### Whole genome sequencing, assembly and annotation

Bacterial genomic DNA was extracted using ZR Fungal/Bacterial DNA MiniPrep Kit (Zymo Research, Irvine, CA, USA) and quantified using Qubit 2.0 Fluorometer (Thermo Fisher Scientific, Waltham, MA, USA). 1 ng of DNA sample was used in the preparation of Illumina sequencing libraries using Nextera XT sample preparation kit with dual indexing following provider’s instructions. Sequencing libraries were pooled and sequenced in-house on Illumina MiSeq platform with 2*250 bp pair-end sequencing kit.

The raw sequencing reads were assembled into the high-quality draft genome using SPAdes v3.10 (Bankevich, Nurk et al. 2012) which is a de Bruijn graph-based assembler for the bacterial genome. Quality of the assembled genome was accessed using QUAST v4.4 (Gurevich, Saveliev et al. 2013) and overall coverage of assembled genome was calculated using BBMap (Bushnell 2014). The assembled genomes were annotated using NCBI prokaryotic genome annotation pipeline (Tatusova, DiCuccio et al. 2016).

### Whole genome based phylogeny

Phylogenomic analysis based on more than 400 putative conservative genes was carried out using PhyloPhlAn v0.99 (Segata, Börnigen et al. 2013). Here, USEARCH v5.2.32 (Edgar 2010), MUSCLE v3.8.3 (Edgar 2004) and FastTree v2.1 (Price, Dehal et al. 2009) were utilized for orthologs searching, multiple sequence alignment and phylogenetic construction respectively. All strains of XSXP phylogroup and *Luteimonas mephitis* DSM12574^T^ (used as an outgroup) were used for construction the phylogeny.

In order to obtain a more robust whole genome phylogeny, core genome based phylogeny was constructed using MAFFT v7.31 (https://mafft.cbrc.jp/alignment/software/) and the FastTree v2.1 (Price, Dehal et al. 2009) which was integrated within the roary v 3.11.2 (Page, Cummins et al. 2015) with identity cut-off of 60%. Implementation of the tools can be found in detail under the section of pan genome analysis.

### Taxonogenomics analysis

Genome relatedness was assessed using the average amino acid identity (AAI), percentage of conserved protein (POCP) and average nucleotide identity using (ANI). AAI was calculated using CompareM v0.0.23 (https://github.com/dparks1134/CompareM), which uses the mean amino acid identity of orthologous genes between a given pair of genomes. POCP is another method to evaluate the genome relatedness at genus level, which is based on amino acid conservation. POCP is calculated with the blast search using the default settings (Qin, Xie et al. 2014) (https://figshare.com/articles/POCP-matrix_calculation_for_a_number_of_genomes/5602957). Further, fastANI v1.3 (Jain, Rodriguez-R et al. 2018) an alignment-free sequence mapping method with default settings was used to calculate ANI values with default settings.

### Cluster analysis

Protein sequences of the gene clusters (Bansal, Midha et al. 2019) were used as query and tBLASTn searches were performed on the XSXP phylogroup genomes. Here, tBLASTN searches were performed using standalone BLAST+ v2.9.0 and cut-offs used for identity and coverage were 50% and 50% respectively. Heatmap for the blast searches were generated using GENE-E v3.03215 (https://software.broadinstitute.org/GENE-E/).

### Pan genome analysis

Pangenome analysis of the strains was performed using Roary v3.12.0 (Page, Cummins et al. 2015). Here, .gff files generated by Prokka v1.13.3 (Seemann 2014) were used as input for Roary pan genome analysis. Since, we are analyzing genomes from different species, identity cutoff used was 60%. Functional annotation of the core genes identified was performed using EggNOG v4.5.1 (Jensen, Julien et al. 2007).

## Author Contributions

SK, SS and PP has carried out strain procurement from culture collection and strain revival. SK has performed whole genome sequencing and submission of assembled genome to NCBI. KB, SK and AK carried out all the bioinformatics analysis. KB and PBP drafted the manuscript with input from all the authors. PBP has conceived the study and participated in the design. All the authors have read and approved the manuscripts.

## Conflict of Interest Statement

The authors declare that the research was conducted in the absence of any commercial or financial relationships that could be construed as a potential conflict of interest.

## Acknowledgments

This work was supported by a project entitled “Mega-genomic insights into coevolution of rice and its microbiome” (MLP0020) of Council of Scientific and Industrial Research (CSIR) to PBP.

## Supplementary material

**Supplementary figure 1: Heatmap showing average nucleotide identity (ANI) for XSXP.** Dendogram based on ANI values is also shown.

**Supplementary table 1:** Genes unique to Xcc

**Supplementary table 2:** Genes unique to Xhc

**Supplementary table 3:** Genes unique to Xfc

**Supplementary table 4:** Genes unique to Smc

