## Supplementary figures and images for "Deep phylo-taxono-genomics (DEEPT genomics) reveals misclassification of *Xanthomonas* species complexes into *Xylella, Stenotrophomonas* and *Pseudoxanthomonas*"

### supplementary figure 1

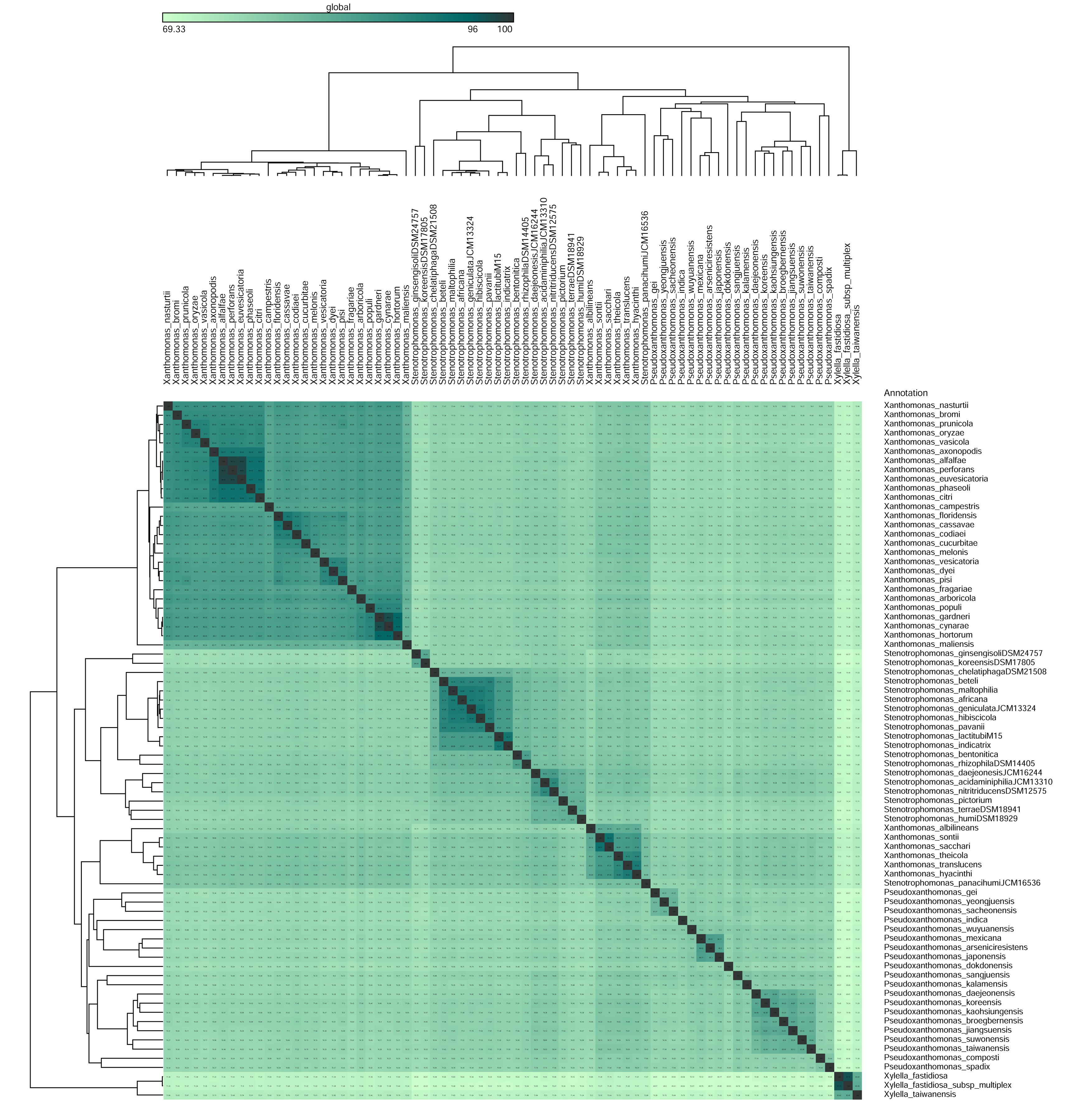
